# Genetic, lifestyle and environmental risk factors for chronic pain revealed through GWAS

**DOI:** 10.1101/2020.05.26.115568

**Authors:** Mischa Lundberg, Adrian I. Campos, Scott F. Farrell, Geng Wang, Michele Sterling, Miguel E. Renteria, Trung Thanh Ngo, Gabriel Cuellar-Partida

## Abstract

Chronic pain (CP) is a leading cause of disability worldwide with complex aetiologies that remain elusive. Here we addressed this issue by performing a GWAS on a large UK Biobank sample (N=188,352 cases & N=69,627 controls) which identified two independent loci associated with CP near *ADAMTS6* and *LEMD2*. Gene-based tests revealed additional CP-associated genes (*DCAKD, NMT1, MLN, IP6K3*). Across 1328 complex traits, 548 (41%) were genetically correlated with CP, of which 175 (13%) showed genetic causal relationships using the latent causal variable approach and Mendelian randomization. In particular, major depressive disorder, anxiety, smoking, body fat & BMI were found to increase the risk of CP, whereas diet, walking for pleasure & higher educational attainment were associated with a reduced risk (i.e., protective effect). This data-driven hypothesis-free approach has uncovered several specific risk factors that warrant further examination in longitudinal trials to help deliver effective early screening & management strategies for CP.

## Introduction

Chronic pain (CP) is a leading cause of disability affecting around 30%–50% of the world’s population with enormous costs to society [6,23,26,34,49,68,77,86]. CP is typically defined as pain that persists or recurs for more than three months and is considered a clinical entity in its own right [65,71,81]. Although CP often arises as a result of injury or disease, not everyone who is injured will develop the condition. This observation suggests that variation in and interactions between individuals’ environments, lifestyles and genetic makeup play a critical yet elusive role in the aetiology of CP.

Epidemiological studies of CP have identified a myriad of risk factors such as sex and age (demographics), diet and alcohol consumption (lifestyle), and clinical comorbidities (e.g., psychiatric and chronic cardiometabolic conditions) [54]. Evidence for the role of genetic factors in CP comes from twin and family studies [38] where its heritability is estimated to be as high as 60% [11,37]. More recently, large-scale genome-wide association studies (GWAS) have identified multiple genetic variants associated with chronic back pain [25,75] and multisite chronic pain [36] as further evidence of a genetic contribution to CP. However, their causal relationship has yet to be adequately examined. Given the complex multifactorial nature of CP, determining its aetiology in large population samples would not be feasible through conventional experimental methodologies.

In this study, we investigated the genetic aetiology of CP by conducting a GWAS using UK Biobank data [57]. We then used these results in a comprehensive set of downstream analyses to draw insights into the environmental, lifestyle and clinical factors linked to CP. These analyses included estimating the genetic correlation of CP with > 1,300 complex traits and conditions, followed by an assessment of whether the genetic correlations could be explained by a causal relationship through applying Mendelian randomization [90] and latent causal variable analysis [59].

## Methods

### Sample selection

Data for CP was obtained from UK Biobank participant responses to the questions: “Have you had back pains for more than 3 months?”, “Have you had hip pains for more than 3 months?”, “Have you had knee pains for more than 3 months?”, “Have you had neck or shoulder pains for more than 3 months?”, “Have you had abdominal pains for more than 3 months?”, “Have you had facial pains for more than 3 months?”, “Have you had headaches for more than 3 months?”, “Have you had pains all over the body for more than 3 months?” (Questionnaire field ID: 6159) [57]. Each of these questions could be answered with “Yes”, “No”, “Don’t know”, or “Prefer not to answer”. Individuals that selected “Yes” to any of the questions were defined as cases while those that selected “No” to all the questions were used as controls. The count of individuals included in the analysis reporting pain for more than 3 months and in which body site is shown Table *1*. In total, 188,352 (73%) individuals that reported pain at any site for over 3 months (cases) while 69,627 reported not experiencing any pain for over 3 months (controls) were included in the analysis. Informed consent was provided by all participants. The research was approved by the UK Biobank’s governing Research Ethics Committee.

### Genome-wide association study

Association testing between CP and genetic variants was conducted using a linear mixed model implemented in Bolt-LMM (v. 2.3.4) [47]. Quality control included exclusion of variants with minor allele frequency (MAF) < 0.005, imputation quality < 0.6 and those deviating from Hardy–Weinberg equilibrium (P-value < 1×10^−5^). Subject data were excluded if their genotype-derived principal components 1 and 2 were further than 6 standard deviations away from those of 1000 Genomes European population. Sex, age and the top 20 principal components derived from genetic data were used as covariates for the association analysis. In addition, a sex-stratified GWA analysis of CP was performed using the same parameters but without sex as a covariate.

### Post-GWAS analyses

Post-GWAS analyses were performed on the Complex Traits Genetics Virtual Lab (CTG-VL) [18] — a freely available web platform for running commonly used statistical genetics analysis methods alongside GWAS summary-level statistics of 1,328 traits & diseases. As described further below, these downstream analyses used summary statistics from the CP GWAS to carry out gene-based and gene-set enrichment analyses, genetic correlation estimates and assess their causal relationship.

### Gene-based & gene-set enrichment analyses

Gene-based association analysis was carried out using the fastBAT [3] implemented in CTG-VL [18]. fastBAT estimates the overall association of SNPs within a 50kb from each gene using GWAS summary statistics and reference genotypes from the 1000 Genomes project European population to account for linkage-disequilibrium between the SNPs. Through this method, we tested the association of 24,254 genes with CP. The Bonferroni’s corrected P-value level of significance was 2.05×10^−6^.

Gene-set enrichment analyses were performed using DEPICT [63] (v. 1 rel. 194) which is able to identify pathways and tissues/cell types that are most likely to underlie GWAS associations. Briefly, DEPICT assesses whether genes in associated loci are highly expressed in any of the 209 Medical Subject Heading (MeSH) tissue and cell type annotations based on RNA-seq data from the GTEx project [78]. Further, it assesses whether there is an enrichment of genes identified in particular molecular pathways constructed based on 14,461 gene sets from diverse databases [63]. As input for DEPICT, we used independent SNPs (based on clumping with an *r*^2^ (LD) between SNPs < 0.05 and 1Mb windows) with a P-value < 1×10^−5^.

### Heritability, Genetic correlations & causality analyses

Genome-wide SNP Heritability 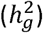 was calculated using LDSC. We transformed the *h*^2^ estimate from observed to liability scale using prevalence estimates of our sample (73%) and a population prevalence of 50 % [12,23,30,51,53,75,85,86,91,92] using the commands --samp-prev 0.73 and -- pop-prev 0.5. The liability scale heritability 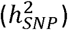 corrects for ascertainment bias regarding the prevalence of the condition.

We estimated pair-wise genome-wide genetic correlation (r_G_) between CP and 1,328 phenotypes using Linkage Disequilibrium Score Regression (LDSC) [9,10] implemented in CTG-VL [18]. To assess whether statistically significant (FDR < 5%) genetic correlations observed could be due to a causal relationship between traits, we then applied the Latent Causal Variable (LCV) method [59]. Briefly, LCV estimates the genetic causal proportion (GCP) whose absolute value ranges from 0 (no partial genetic causality) to 1 (full genetic causality). A high GCP value indicates that trait one is likely to affect trait two. A negative value indicates that trait two affects trait one. In our study CP was set as trait 1, meaning that a positive GCP suggest that CP affects the tested trait while a negative GCP indicates that CP is affected by the tested trait.

Generalized Summary-data-based Mendelian Randomization (GSMR) [90] implemented in gcta (v. 1.92.4beta) was used in addition to the LCV method to assess for potential causal relationships. For this, we used GWAS summary statistics for those traits displaying significant genetic correlations and that had a least three independent instrumental variables (i.e. SNPs with an association P-value < 5×10^−8^ with the exposure). Furthermore, given that two-sample MR may be biased due to weak instruments when both, the exposure and the outcome GWAS are drawn from the same sample, as a sensitivity analysis, we leveraged the sex-stratified GWAS of CP and the sex-stratified GWAS summary statistics from Neale lab [57] [http://www.nealelab.is/uk-biobank] to perform GSMR analysis using GWAS summary statistics of CP in males and GWAS summary statistics of exposures in females and vice versa, thus ensuring non-overlapping samples.

## Results

### Genome-wide association of chronic pain

After quality control (Methods), our GWAS included 188,352 (73%) CP cases (individuals that reported pain at any site for over 3 months) and 69,627 controls. No evidence for inflated statistics due to hidden population stratification was detected (the LDSC intercept estimate was 1.002 ± 0.008). The SNP heritability of CP on the liability scale was 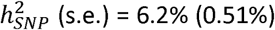. Two loci were identified at genome-wide levels of significance (P-value < 5×10^−8^) (Figure 1 and Table *2*). The lead SNPs at these loci were rs10660361 and rs113313884 which are located in the intronic region of *LEMD2* in chromosome 5 and *ADAMTS6* in chromosome 6 respectively (Figure 1b and 1c). Using Complex Traits Genetics Virtual Lab (CTG-VL) [18], we explored whether those SNPs have been previously associated to other traits. We observed that rs10660361 was also identified by a recent GWAS on back pain (P-value = 7.77×10^−8^) [25] and multisite chronic pain (P-value = 1.9×10^−9^) [36]. However, rs113313884 was only associated with multisite chronic pain at a suggestive level (P-value = 6.9×10^−3^) highlighting the importance of studying phenotypes using different definitions. We observed that these SNPs are also associated to anthropometric traits including height, BMI, and body fat percentage. Furthermore, rs10660361 is strongly associated to platelet count (P-value = 4.19×10^−60^). Results for all significant and suggestive associations are included in Supplementary Table 1.

**Figure 1.**
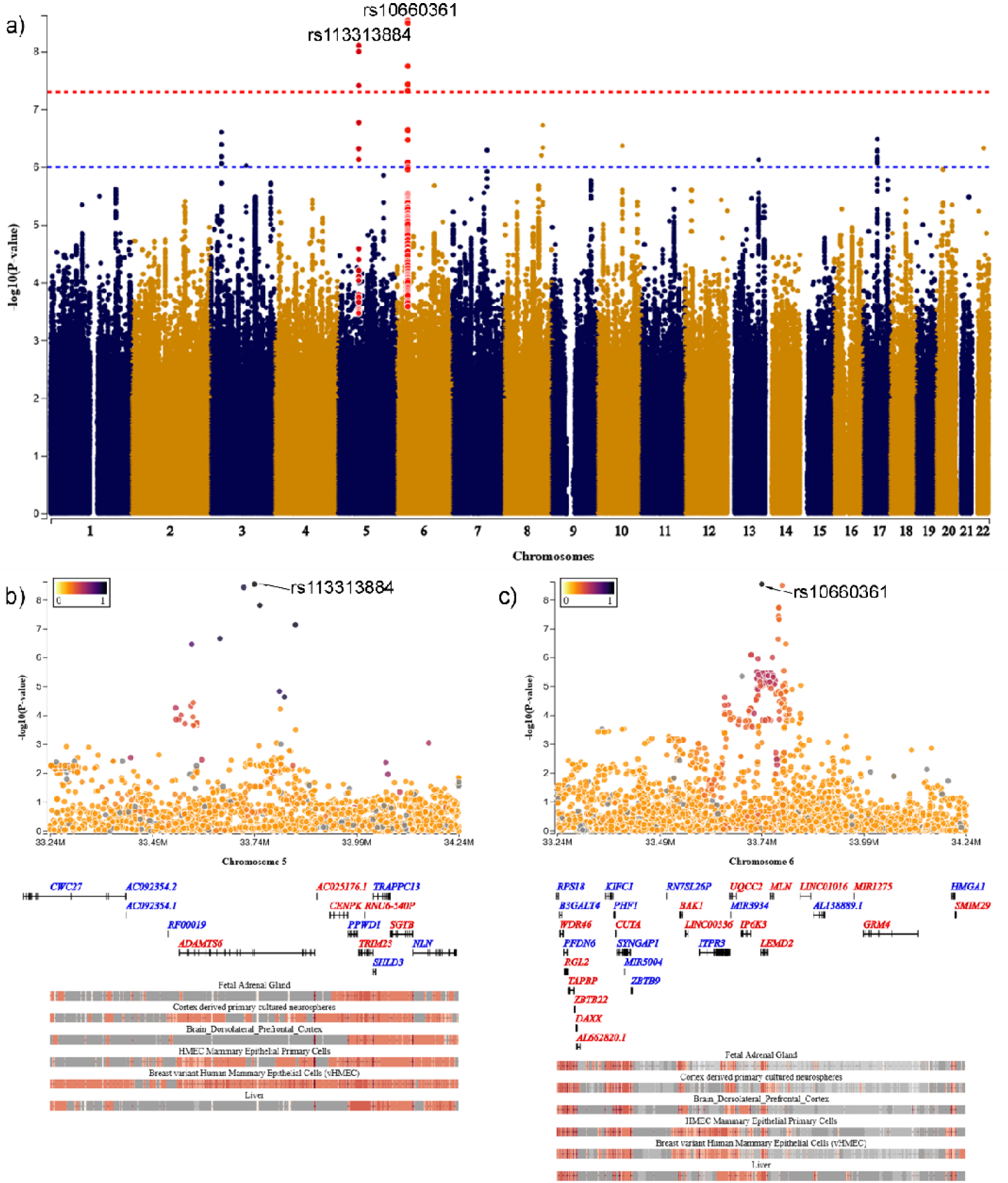
(a) Manhattan plot of CP GWAS showing two significant loci after multiple testing, rs10660361 and rs113313884 mapping to the genes LEMD2 and ADAMTS6 respectively. Annotated are the top SNPs of the discovery GWAS. (b) and (c) LocusTracks of main GWAS. LocusTrack plot for rs113313884 (b) and rs10660361 (c) displaying chromatin state annotation tracks for specific tissues. Red coloured positions in the annotation track indicate open chromatin/active transcription. Tissue tracks in the plots were predefined through DEPICT tissue enrichment analysis top results (see Supplementary Table 3 and 4) of CP.

Epidemiological studies have established a difference in prevalence of CP between sexes (∼60% females and ∼40% males by age 50) [1,4,8,21,61,64,72,73]. In line with this, data from UK Biobank showed a prevalence of 56.76% in females and 43.24% in males. Here, we sought to explore whether this difference in prevalence may be partly due to genetic differences (e.g. genes involved in hormone regulation may play a role in CP) by performing a sex-stratified GWAS of CP. There were no loci associated with CP in the sex-stratified GWAS at genome-wide levels of significance (P-value < 5×10^−8^). Nevertheless, the association of *LEMD2* (rs10660361) and *ADAMTS6* (rs113313884) with CP was still suggestive in both, males (P-value = 1.9×10^−7^, Beta (s.e.) = 1.1×10^−2^ (2.0×10^−3^) and P-value = 5.1×10^−5^, Beta (s.e.) = 2.4×10^−2^ (5.8×10^−3^)) and females (P-value = 1.3×10^−3^, Beta (s.e.) = 5.5×10^−3^ (1.7×10^−3^) and P-value = 2.8×10^−5^, Beta (s.e.) = 2.0×10^−2^ (4.9×10^−3^)). The genetic correlation derived from a bivariate *Linkage Disequilibrium Score Regression* (LDSC) [10] analysis between GWAS of CP in males and CP in females suggested that CP in males and females is influenced by the same genetic factors r_G_ (s.e.) = 1.0 (0.1022).

### Gene-based, pathway and tissue enrichment analyses of chronic pain

The gene-based association analysis revealed five significant (P-value < 2.06×10^−6^) genes associated with CP (Table *2*). The association of three of the genes (*LEMD2, MLN, and IP6K3*) was driven by the lead SNP rs10660361 identified in the GWAS. However, *DCAKD* and *NMT1* were in the 17q21.31 locus that was not identified during the main GWAS. Full results of the *fast and flexible gene- or set-Based Association Test* (fastBAT) [3] analysis are shown in Supplementary Table 2. Pathway and tissue enrichment analyses performed using DEPICT [63] did not identify any pathways or tissues associated with CP (false discovery rate < 5%). The full results for pathway and tissue enrichment analyses are shown in Supplementary Table 3 and 4.

**Table 1.**
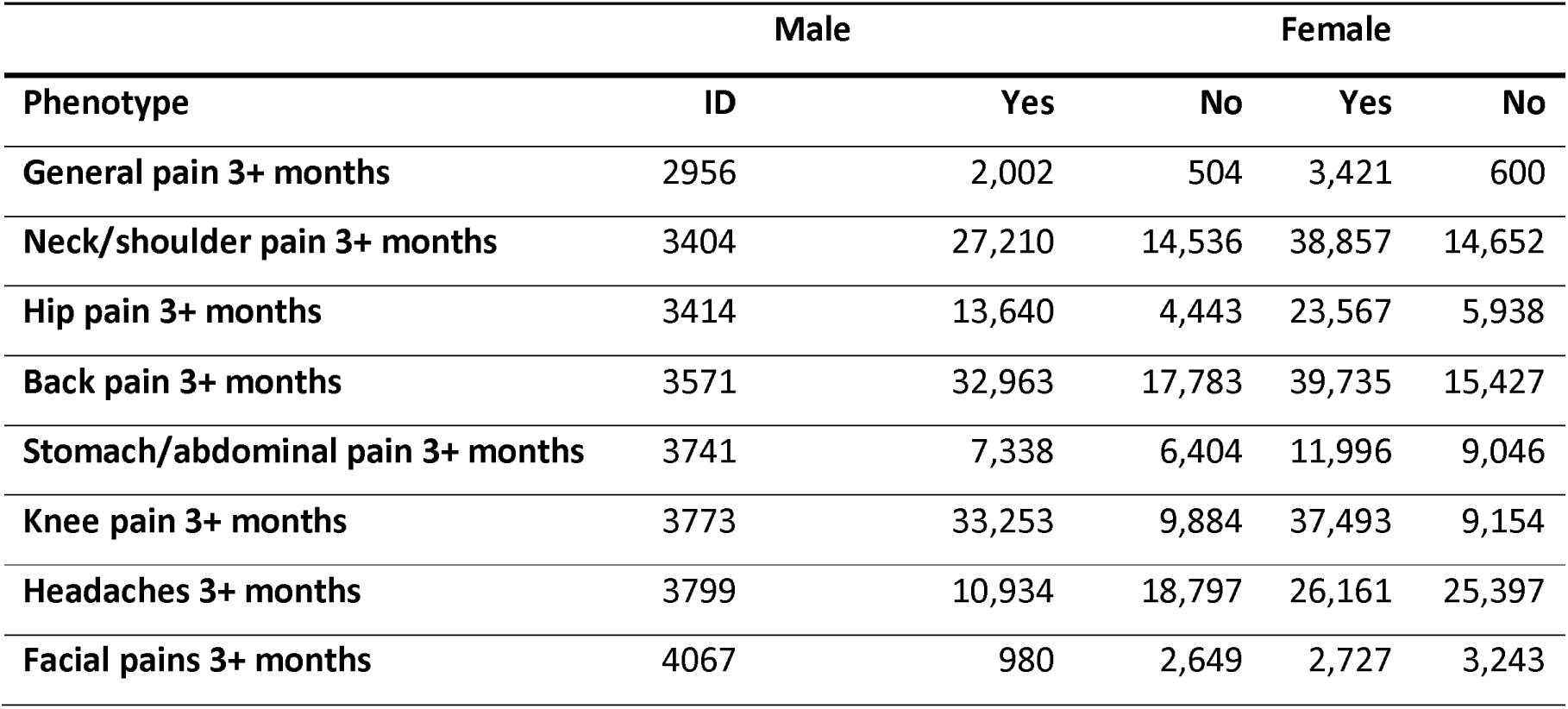
Phenotypes included from UK Biobank to define chronic pain. If an individual answered yes to having pain in any of the sites below, it was considered a chronic pain case. Only individuals of European descent were included in the analysis.

**Table 2.** Lead SNPs identified in discovery GWAS and significant results (P-value < 2.06×10^−6^) of the fastBAT gene-based test. Genomic positions are shown in genome assembly GRCh37 (hg19).

| SNP | CHR:POS | Gene closest to<br>SNP | A1/A2 | A1<br>FREQ | OR (s.e.) | P-value |
| --- | --- | --- | --- | --- | --- | --- |
| **rs10660361** | 6:33741371 | *LEMD2* | C/CG | 0.5756 | 1.04<br>($6.59 \times 10^{-3}$) | $2.90 \times 10^{-9}$ |
| **rs113313884** | 5:64629986 | *ADAMTS6* | C/T | 0.9723 | 1.12<br>(0.0187) | $7.90 \times 10^{-9}$ |

fastBAT gene-based test results
| Gene Name | CHR | Start | P-value | Top SNP | #SNPs |
| --- | --- | --- | --- | --- | --- |
| ***LEMD2*** | 6 | 33738989 | $1.97 \times 10^{-8}$ | rs10660361 | 170 |
| ***MLN*** | 6 | 33762448 | $7.46 \times 10^{-8}$ | rs10660361 | 185 |
| ***DCAKD*** | 17 | 43100705 | $1.01 \times 10^{-6}$ | rs2301597 | 144 |
| ***NMT1*** | 17 | 43138679 | $1.14 \times 10^{-6}$ | rs2301597 | 152 |
| ***IP6K3*** | 6 | 33689442 | $1.75 \times 10^{-6}$ | rs10660361 | 136 |

Allele 1 (A1) represents the effect allele (which increases risk for the tested trait) is and Allele 2 (A2) represents the non-effect allele. Beta describes the effect of this SNP with CP and its direction. MAF is the minor allele frequency. OR is the odds ratio and thus, is a ratio between the odds for having the effect allele and the odds of not having the effect allele.

### Genetic correlations and causality assessment

Bivariate LDSC analyses [9,10] between CP and 1,328 traits revealed 548 genetically correlated traits at Benjamini-Hochberg’s *false discovery rate* (FDR) < 5% (Supplementary Table 5). As expected, the largest positive genetic correlations were observed for pain-related traits in addition to musculoskeletal conditions. We also observed positive genetic correlation with sleep disorders, including sleep apnoea and psychiatric traits and disorders including anxiety, attention deficit and hyperactivity disorder (ADHD) as well as depression. Current smoking had also a positive genetic correlation with CP while alcohol intake variables such as being a current alcohol consumer of whether alcohol is taken during meals was negatively correlated with CP.

To assess whether the observed genetic correlations observed were due to pleiotropy or a causal relationship, we used the *Latent Causal Variable* (LCV) approach [59] to estimate the *genetic causal proportion* (GCP) on the 548 traits displaying genetic correlations at FDR < 5%. The causal architecture plot [29] summarizes the LCV results is shown in Figure 2. Supplementary Table 5 shows the complete results for these analyses. Briefly, traits displaying a negative GCP suggest that they may cause CP, while traits with a positive GCP suggest that the trait may be caused by CP. Furthermore, the sign of the genetic correlation coefficient indicates whether the causal trait decreases or increases the risk of CP. In total, we identified 84 potential causal relationships at FDR < 5% using the LCV method, most of them (83/84) affecting CP rather than CP having an impact on them. Among traits with evidence of a causal relationship, we found that dietary behaviours such as consumption of cereal, starch and tea were associated with a reduced risk of CP while vitamin supplementation was associated with an increased risk of CP. These results may arise due to underlying conditions that lead to the intake of vitamins and CP. As expected, we observed that fibromyalgia related traits and musculoskeletal conditions including arthritis and spondylitis had the strongest evidence of causally increasing the risk of CP. Furthermore, we found that endocrine, nutritional and metabolic diseases including having high cholesterol may increase the risk of CP. Finally, there was some evidence of major depressive disorder and sleep disorders increasing risk of CP. Nevertheless, both associations were below the FDR threshold.

**Figure 2.**
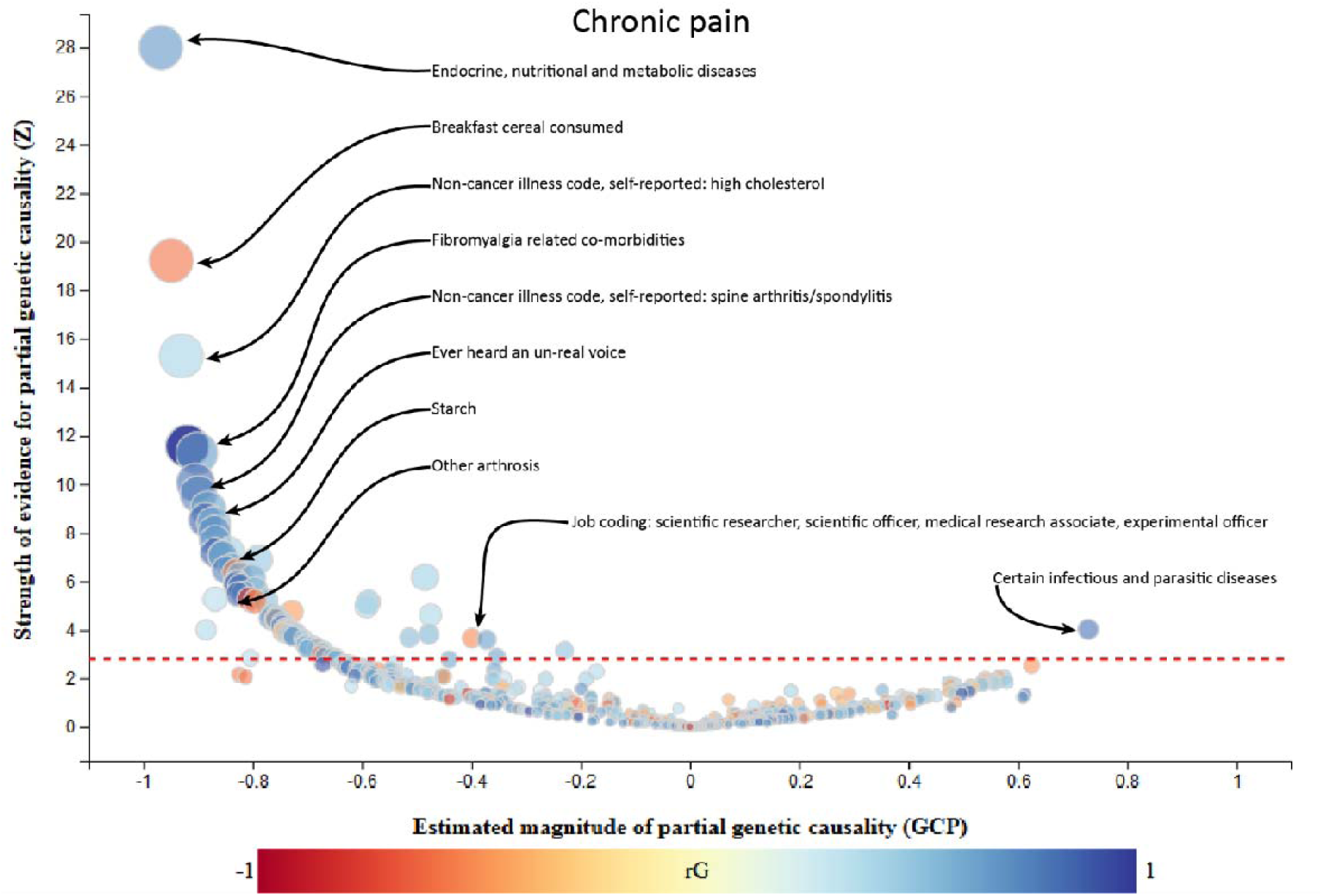
LCV results of CP vs 548 traits displaying genetic correlations. On the X-Axis the likelihood of CP affecting a trait (+1) or the trait affecting CP (−1) is represented. The Y-Axis illustrates the strength of evidence of a causal association (Z score for H_0_: GCP = 0). The colour indicates the genetic correlation, which ranges from −1 to 1 and indicates whether the trait increases or decreases the risk of CP.

To complement the LCV analysis, we conducted Mendelian randomization using the Generalized Summary-based Mendelian Randomization (GSMR) [90] approach for all the traits displaying genetic correlations. Using GSMR, we were unable to assess if CP affected any trait, as there were insufficient instrumental variables to proxy CP. However, we could identify multiple factors affecting CP. This included evidence of higher body mass index (BMI), whole-body fat mass, waist hip ratio, among other body composition traits to increase the risk of CP. Diseases of the musculoskeletal system and connective tissue including arthrosis displayed an increase of risk of CP. Finally, we also found evidence of educational attainment reducing the risk of CP and Major Depressive Disorder (MDD), mood swings, ADHD, anxiety and smoking increasing the risk of CP.

Given that GSMR can be susceptible to bias (e.g. weak instrument bias) when the exposure and the outcome (CP) GWAS are drawn from the same samples, we leveraged the sex-stratified CP GWAS results, in addition as sex-stratified GWAS summary data from Neale’s Lab (see Methods section) [83]. Briefly, we used GWAS performed using only female samples as exposure and GWAS estimated using male samples as outcome (CP) and vice versa. Consequently, we were able to confirm that an increase in BMI, body fat and waist-hip ratio as well as major depressive disorder and anxiety increase the risk of CP while education reduces risk. All results are shown in Supplementary Table 5.

## Discussion

Here we present one of the largest GWAS of CP. We leveraged summary-based data to identify factors that may increase or reduce risk of CP, including genetic, lifestyle and clinical. We discovered variants near *ADAMTS6* and *LEMD2* associated with CP during our main GWAS analysis. *ADAMTS6* has been reported to show increased levels of expression in patients with osteoarthritis [69,76,88] and is associated with heel bone mineral density, which is used in diagnosis of osteoporosis [40,55]. The SNP rs10660361 was closest to *LEMD2*, a gene implicated in embryonic development and the regulation of several signalling pathways [46]. However, we note that this locus was reported in a previous GWAS on multisite chronic pain mapping to the gene *MLN* [36]. It is important to note that we were unable to map the genes likely affected by these SNPs through more robust approaches (rather than only based on proximity) such as eQTLs. Our tissue enrichment analysis was unable to identify putative cells/tissues of interest where we could look for potential eQTLs. A lookup in the GTEx eQTL database [78] only show suggestive evidence of rs113313884 being associated to the expression of *TRAPPC13* in tibial arteries.

Our gene-based analysis implicated the 17q21.31 locus, particularly the genes *DCAKD* and *NMT1* to be associated with CP. This locus also harbours an inversion polymorphism that has been implicated in numerous neurodegenerative diseases and cognition [80]. *DCAKD* has previously been associated to psychiatric disorders including schizophrenia and bipolar disorder [67]. These results suggest that the association of CP with psychiatric disorders may be partially driven by genetic variation at this locus. Furthermore, *DCAKD* has also been associated with osteoarthritis which is also a potential cause of pain [39]. *NMT1* was already reported to be associated to multisite chronic pain [36], in addition to pulse pressure and insomnia [27,36]. Besides *DCAKD* and *NMT1*, the fastBAT gene-based test also revealed significant associations with *MLN* and *IP6K3*. Both, *MLN* and *IP6K3* are mapped to rs10660361. Polymorphic variations of *IP6K3* contribute to the susceptibility of late onset Alzheimer’s disease [17] and BMI [87]. Different variants in *MLN* (*Motilin*) are known for a variety of effects including BMI [15] or depressive symptoms [5]. Further efforts could leverage these results to assess *in vitro* and *in vivo* whether any of these genes is causal for CP, thus identifying potential new therapeutic targets.

The genetic correlation between CP in males and CP in females was not different from the unity (r_G_ = 1.0), suggesting that male and female patients are largely exposed to the same genetic influences and that differences in prevalence may be due to differences in exposure to environmental and lifestyle factors such as smoking, and physical activity amongst others [16,28,41,43]. We identified genetic correlations between CP and a vast array of complex traits ranging from anthropometric to behavioural as well as complex diseases including musculoskeletal and neuropsychiatric disorders. Given that pain is a symptom of many diseases, we carried out a thorough analysis to explore if the genetic correlations were due to horizontal pleiotropy (i.e. the effect of genes on CP is independent from their effect on other traits) or due to a causal relationship.

In line with a recent GWAS on multisite chronic pain [36] and other studies [58,82], we found that depression (including major depressive disorder) and anxiety may increase the risk of CP. This finding is important as so far, studies have been conflicting with regard to whether depression increase the risk of pain or vice versa [13,24,32,35,70,82]. As reported by some epidemiological studies [31,33,48,56,62,74,84] we also observed that increased body fat, BMI and obesity may lead to an increase in the risk of CP. It is hypothesized that the relationship between obesity and pain may be due dysregulation of inflammatory markers due to inflammatory responses [22,52,60]. We also found that traits describing dietary behaviours such as “Starch” consumption appeared to lead to a decreased risk of CP. This was surprising as starch consumption can increase inflammation [19,42,45,50,89]. “Breakfast cereal consumed” also appeared to have a protective effect on CP. It has been documented that dietary cereal fibre lowers risk of osteoarthritis, which is the most common form of arthritis and a leading cause of chronic pain [20]. In addition to dietary fibre, whole grain cereals contain bioactive components including polyphenols, and flavonoids that have anti-inflammatory properties [2,7]. The identified effect of vitamins increasing the risk for CP might be explained by an indirect causal effect. Briefly, an unobserved trait (such as a disease) that increases the likelihood of both vitamin intake and CP could create an apparent causal effect between vitamin supplementation and increased CP, which is confounded by the unobserved trait. In line with this hypothesis, we also observed variables related to medication usage to increase the risk on CP. For example, the trait “Treatment/medication code: dihydrocodeine” (an opioid analgesic) was associated with an increase of risk for CP. While this might be explained by observations linking opioid intake to increased sensitivity of pain [14,44,66,79], it is possible that taking dihydrocodeine can proxy a pain-causing condition.

Our LCV analyses indicated that CP might be a downstream consequence of multiple traits and diseases and is likely not a cause for other conditions. We identified only one trait that might be causally affected by CP. Unfortunately, we were limited to confirm this pattern through Mendelian randomization as our GWAS of CP did not have enough statistical power and genetic instruments to carry out a bidirectional Mendelian randomization analysis, and thus assess the impact of CP on other traits. Another important limitation is that although we found evidence of effects and their direction of several traits and diseases on CP, it is hard to interpret the magnitude of these effects. The genetic causal proportion estimated through the LCV approach only indicates the proportion of genetic correlation that may be explained through a causal relationship, and thus, it does not produce an odds ratio or effect size estimate. The effect estimates of our Mendelian randomization analyses are similarly hard to interpret as GWAS for CP and the majority of diseases tested have been derived through a linear-mixed model, not a logistic regression and thus working out precise effect estimates is challenging.

In conclusion, we have carried out one of the largest and most extensive studies on CP. We highlight three loci that play a role in the risk of developing CP. Identified no evidence of sex-specific genetic effects and provide evidence of a potential causal role for dozens of traits in the development of CP. Our results call for future studies to assess the roles of ADAMTS6, LEMD2, DCAKD, MLN, NMT1, and IP6K3 on CP in vitro and in vivo. We furthermore postulate potential modifiable lifestyle factors related to CP which need to be thoroughly assessed and replicated.

## Acknowledgements

This research was conducted using data from the UK Biobank resource (application number 25331). ML, AIC and GW are supported by a University of Queensland Research Training Scholarship from The University of Queensland (UQ). ML thanks the support of the Commonwealth Scientific and Industrial Research Organisation through a Postgraduate Top-Up Scholarship. MER thanks the support of the NHMRC and Australian Research Council (ARC), through a NHMRC - ARC Dementia Research Development Fellowship (GNT1102821). GC-P is funded by an ARC Discovery Early Career Researcher Award (DE180100976). We thank the members of the Evans Lab and Patricia Gerdes for helpful contributions and discussion. RECOVER Injury Research Centre receives unrestricted grant funding from the Motor Accident Insurance Commission (Queensland). The funders had no role in the design or interpretation of this study.

This research was carried out at the Translational Research Institute, Woolloongabba, QLD 4102, Australia. The Translational Research Institute is supported by a grant from the Australian Government.

## Author contributions

GC-P, TTN and MER conceived and directed the study. AIC and MER performed the GWAS. ML performed downstream statistical and bioinformatics analyses, with support from GC-P. GW, SF, MS and TTN acted as advised in the analyses and/or interpretation of results. ML generated all figures and tables. ML and GC-P wrote the manuscript with feedback from all co-authors.

